# Giant Clam Gardens: Cultural practices and ecological implications for population resilience in New Caledonia

**DOI:** 10.1101/2025.10.12.681949

**Authors:** Pascal Dumas, Cecile Portes, Christophe Peignon

## Abstract

Giant clams (*Tridacninae*) are a key component of coral reef ecosystems and a major economic, subsistence, and cultural resource for Pacific Island communities. In New Caledonia where stocks are declining but giant clams retain an emblematic status, coastal communities have developed a traditional practice referred to as “giant clam gardens,” involving the translocation and aggregation of wild individuals onto shallow reef flats near villages. This study provides the first formal assessment of the prevalence, cultural significance, and potential ecological benefits of this practice. Surveys conducted in 11 coastal tribes documented over 40 clam aggregations primarily composed of *Hippopus hippopus*, which varied greatly in terms of age (1-40 years), area (1-1080 m^2^) and abundance (from a few individuals up to more than 2 700). Our findings suggest that the practice of “giant clam gardens” is now fairly widespread and primarily reflect a pragmatic approach aimed at optimizing resource access, rather than conserving the resource. Population surveys confirmed the presence of five giant clam species (*H. hippopus, Tridacna maxima, T. squamosa, T. derasa* and *T. crocea*), but failed to report significant recruitment in and around aggregations up to 1km downstream from the study area. This work revealed that while notions of protection and conservation were occasionally mentioned during interviews, they were not cited as primary motivations by communities. Nonetheless, clam gardens may contribute to broader population resilience by enhancing reproductive biomass and supporting larval dispersal across larger spatial scales. Given the declining state of natural stocks and the persistence of strong cultural ties to giant clam harvesting, this practice represents a pragmatic, culturally embedded approach that can provide valuable insights for strengthening the sustainability of this resource.

## Introduction

Giant clams are the largest marine mollusks found in coral reef ecosystems of the Indo-Pacific. Known for millennia, they constitute a crucial resource of economic, subsistence, and cultural importance for most island and coastal communities throughout the Pacific (Dalzell et al., 1996; Larson, 2016; Lucas, 1994; Neo and Todd, 2012). These bivalves, particularly the larger species, provide a valuable and easily accessible source of protein and therefore typically support small-scale artisanal fisheries (Mingoa-Licuanan and Gomez, 2002; Raymakers et al., 2003). Their exploitation began shortly after the first human settlements, as evidenced by the numerous artifacts found in the geological strata associated with Lapita populations across the Pacific (Sand, 2010). Certain species, such as *Tridacna maxima, T. crocea* and *T. noae*, are also highly prized in the aquarium trade for their vibrant colors and water filtering abilities (Braley, 2001; Wabnitz, 2003). However, their vulnerability to fishing, combined with quite unfavorable population dynamics (including slow growth and often erratic recruitment), has led to widespread overharvest, resulting in a general decline of most species throughout their distribution range (Bin Othman et al., 2010; CITES, 2004; Neo et al., 2017; Rippe et al., 2024). As a consequence, giant clam species are listed on the IUCN Red List of Threatened Species, with five species currently assessed as Vulnerable or Endangered and one species (*Tridacna gigas*) is Critically Endangered (Neo and Li, 2024); *T. gigas, T. derasa* and look-alike species are protected under Appendix II of the CITES Convention, which regulates the international trade in endangered wild species (Wells, 1997).

In New Caledonia, where they retain an emblematic status and continue to play a role in social and cultural practices, giant clams are primarily harvested for local consumption, often circulating through traditional exchange or sale networks before reaching the final consumer (Virly, 2004). Of the twelve currently recognized species, seven are found across the reef and lagoon environments: *Tridacna maxima, T. crocea, T. squamosa, T. derasa, T. mbalavuana, T. noae*, and *Hippopus hippopus* (Fauvelot et al., 2018; Neo et al., 2017). Their density and distribution differ significantly from one species to another, depending on both their ecological characteristics and their appeal to local fishers (Dumas et al., 2013; Purcell et al., 2020). One species, *Tridacna gigas*, is currently considered extinct in the wild and is only observed in fossil form. The exploitation of giant clams has gradually evolved from subsistence to small-scale commercial fisheries, with the meat being sold in more or less formal markets, either directly or through middlemen. National data on commercial catches indicate substantial but highly variable volumes at the country level, peaking at 9 tons/year in the years 2000-2009. These volumes dropped substantially to around 1 ton/year following the implementation of provincial fishing quotas in 2009. However, the accuracy of these data is limited by the lack of species-specific information, as fishers’ logbooks report all giant clam species collectively. The situation is further complicated by recreational and subsistence fishing activities, generally dominant in the traditional Melanesian context, whose informal and widespread nature makes quantitative estimation particularly difficult (Guillemot et al., 2009; Jimenez et al., 2011). A recent study in coastal communities along New Caledonia’s eastern coast revealed that non-professional catch of fish and invertebrates can exceed professional (i.e. declared) one by 10 to 100 times (Faure et al., 2022). Despite the implementation of protective measures including quotas, size limits and MPAs, fishing pressure is concerning, with local populations of giant clams already showing signs of overexploitation (e.g. reduced densities, smaller individual sizes) in the most frequented lagoon areas (Dumas et al., 2011).

In this context, the creation of “giant clam gardens” is sometimes described as a traditional practice in certain coastal communities (Portes, 2010). These gardens result from the collection of wild giant clams by local fishers during their fishing trips, which are then relocated alive to shallow reef flats near their homes, in areas that are theoretically not accessible to outsiders. Such aggregations have been reported mainly in the Northern Province of New Caledonia, with varying motivations or expected benefits (J. Tiavouane, pers. obs.). In particular, it remains unclear whether they are primarily intended as a means of accumulating biomass (e.g. for household consumption, sale or traditional exchange within the community), and/or as a tool for resource management or conservation. Some fishers might expect their gardens to increase local giant clam stocks through increased reproduction and recruitment within or in the immediate vicinity of the aggregations. However, no quantitative data are currently available in the literature, making it impossible to assess the cultural and ecological significance of this phenomenon. This study therefore aims to: i) assess the existence and prevalence of this phenomenon; ii) gain a better understanding of its cultural significance and the motivations behind the practice, especially in more traditional areas; and iii) quantify clam aggregations in terms of population structure (species abundance, diversity, size distribution) and potential ecological impact on the reef areas where they occur.

## Material & Methods

### 1. Interviews on giant clam knowledge and practices

A preliminary phase was conducted in coordination with the departments responsible for marine resource management in both the Northern and Southern Provinces, with three main objectives: (i) to identify coastal areas potentially hosting clam aggregations; (ii) to establish contact with key informants within the relevant coastal communities; and (iii) to obtain the necessary administrative and customary authorizations (particularly from tribal chiefs) to carry out the field surveys.

Semi-structured, face-to-face interviews were subsequently conducted in the target areas using a standardized questionnaire, in the presence of an authorized province field officer to facilitate communication and, when necessary, provide translation of local dialects (e.g., places or species names). The questionnaire was designed to investigate practices and local knowledge related to giant clams. In addition to collecting background information on respondents, it comprised 40 items divided into two main sections: a) a general section focused on giant clam consumption and harvesting practices; and b) a specific section aimed at characterizing clam aggregations (e.g., location, area covered, age, environmental attributes) and associated practices (e.g., selection criteria, objectives, local management, potential issues…) (Supp. Table 1). Each interview lasted approximately 30-40 minutes.

### 2. Giant clam aggregation surveys

Following the interviews, quantitative field surveys were conducted to assess the characteristics and population structure of the clam aggregations, including the area covered, species abundance, species richness, and size distribution. These surveys were conducted in the coastal area of the Tiari tribe, Northern Province, on the eastern coast of New Caledonia (20°15.042 ′;S, 164°21.204 ′;E), where the largest and most abundant aggregations were reported during the questionnaire phase.

Fieldwork was conducted at low tide in and around Tiari Bay, an open bay encompassing approximately 3 km of coastline located at the extreme north of the east coast of New Caledonia (Fig. 1). For each aggregation, a rectangular perimeter was delineated to enclose the area and allow for surface area estimation. The geographic coordinates of the center of each aggregation were recorded using a Garmin 60 GPS device. Species identification and counting were performed by a team of 2 to 4 individuals working simultaneously. Depending on their size, clam aggregations were subdivided into smaller sections to facilitate individual identification and counting. Within each aggregation, all clam individuals were identified to species level and counted. Additionally, shell length (measured to the nearest 5 mm) was recorded for approximately 20% of randomly selected individuals to assess size structure within each aggregation.

**Figure 1.**
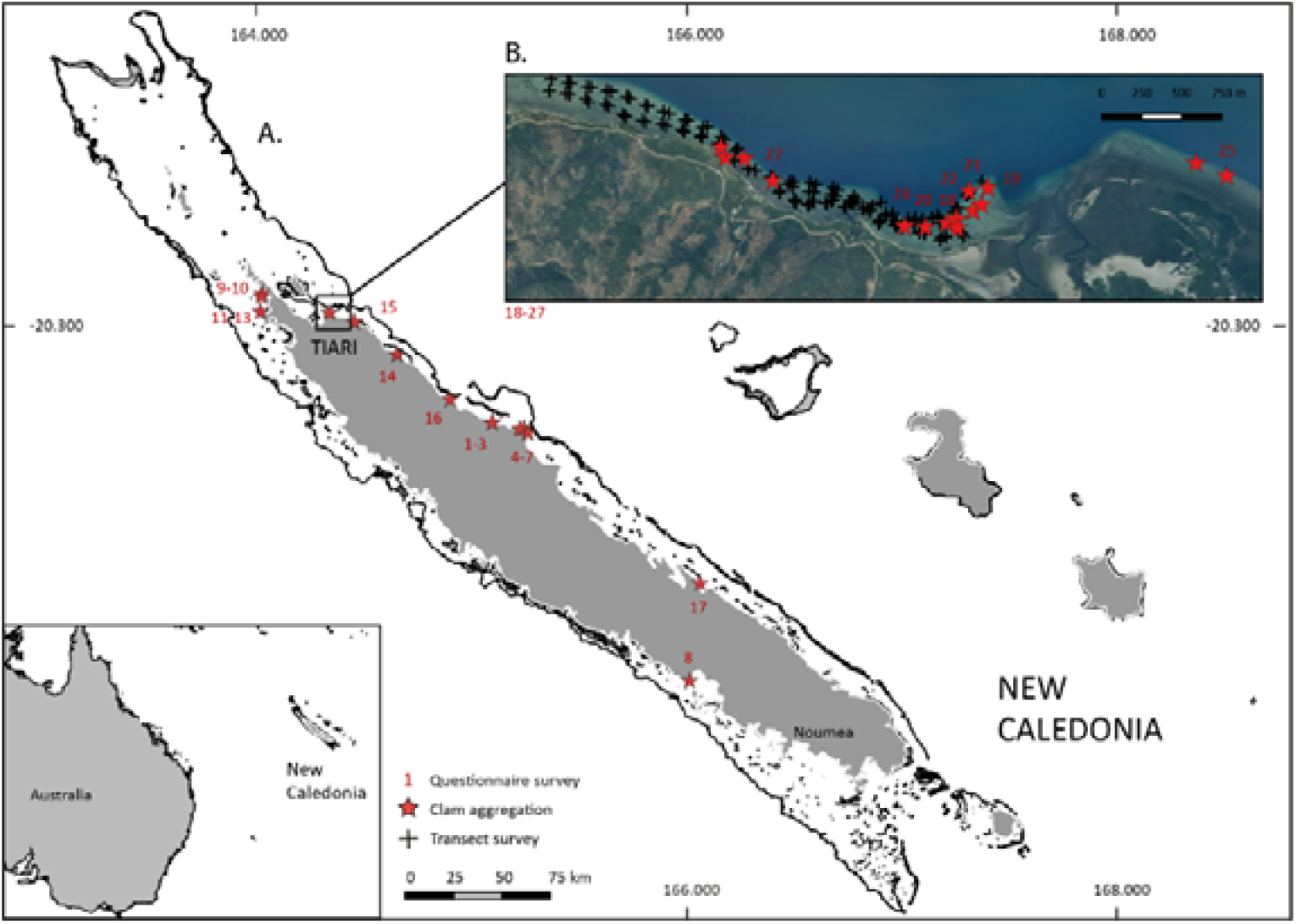
Location of giant clam surveys across New Caledonia, South Pacific. A. Country scale. B. Bay of Tiari (North Province, northeastern coast).

### 3. Giant clam population surveys

The structure of natural giant clam populations, with particular attention to the presence of juveniles near the aggregations, was investigated on the fringing reefs in and around Tiari Bay using underwater transects. Prior to the surveys, geomorphological classifications and available habitat maps were used to confirm that the area featured reef habitats suitable for supporting natural giant clam populations.

Data were collected at high tide by a team of 1 to 4 snorkelers, swimming parallel to the shoreline along 100-meter transects. Surveys began in the southern part of Tiari Bay and extended northward, following the direction of the prevailing currents. The transects were conducted within a depth range of 1–5 meters, with a minimum spacing of 10 meters between adjacent transects. All giant clams that were detectable along a 1 m-wide corridor were identified to the species level and counted. Individual sizes were recorded to the nearest 5 mm using calipers. All transects were georeferenced by marking both endpoints with a handheld Garmin GPSMap 60Cx in an underwater housing.

## Results

### 1. Interviews on giant clam knowledge and practices

Surveys based on information provided by the provincial services allowed us to identify 39 giant clam aggregations, through 27 individual questionnaires conducted in 11 coastal tribes throughout the territory (Figure 1). The vast majority of these aggregations (84.6%) were located on the east coast of New Caledonia, with a smaller number identified on the west coast (15.4%). A particularly high concentration was recorded in the Tiari tribe, located in the far north of the east coast, with 13 clam aggregations reported during interviews. Four additional aggregations were observed in the Tiari area during the subsequent in situ survey, but they were not included in the interviews.

Survey results indicated that most aggregations were established by non-professional fishers (73.1%), predominantly men (74.1%). A single individual could maintain several aggregations (range: 1–4), which varied greatly in terms of area covered (1 - 1080 m^2^) and age (1 - 40 years). The number of clams per aggregation ranged from a few individuals to several hundred, with the largest exceeding 2 000 individuals (Table 1; Figure 2a).

**Figure 2.**
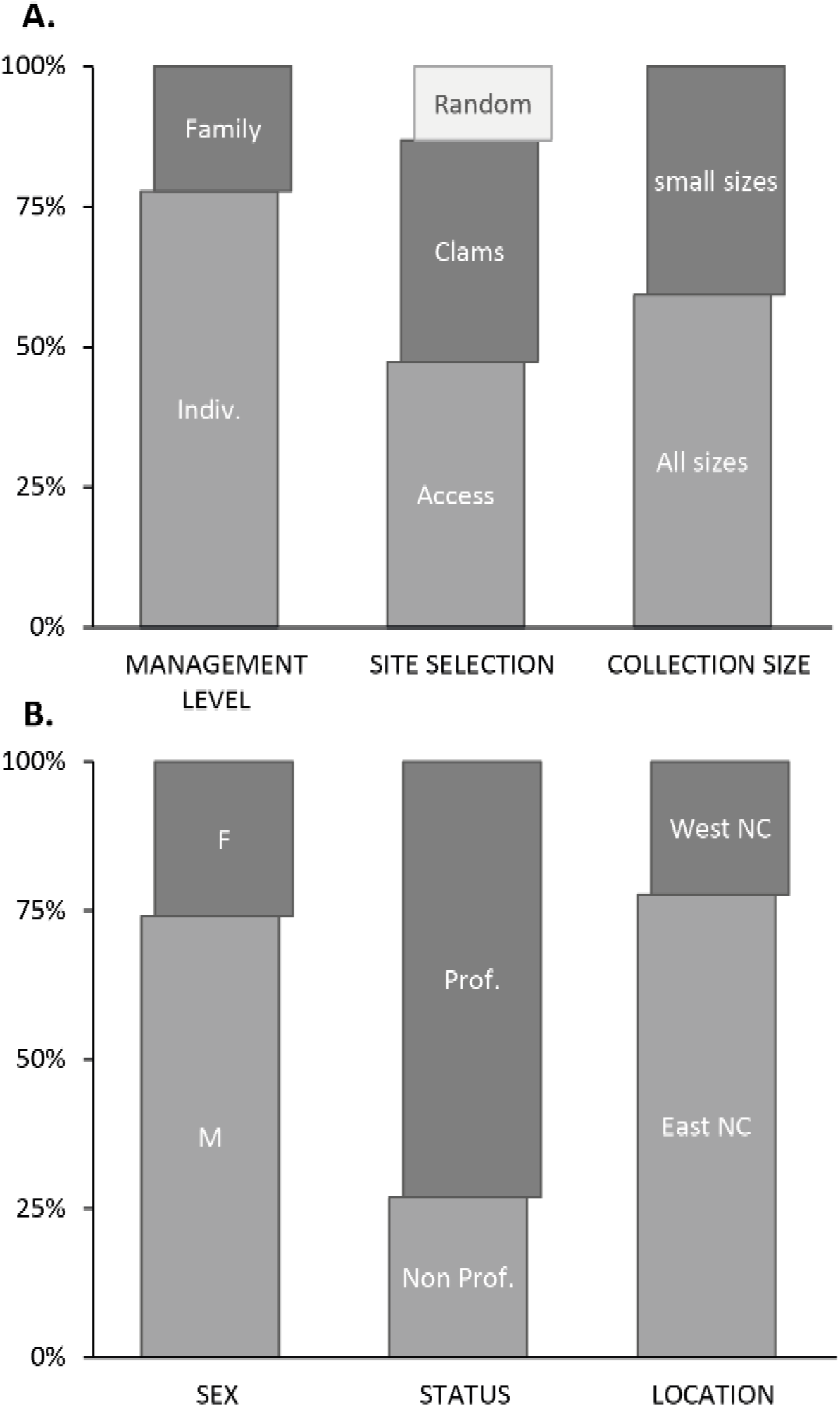
Overview of giant clam aggregation characteristics in New Caledonia, South Pacific. A. Owner: sex, status (professional vs. non-professional fisher), and location (eastern vs. western coast of New Caledonia). B. Aggregation: management level (individual vs. family vs. tribe), rationale for site selection (random vs. existing clam populations vs. ease of access), and size of collection (all sizes vs. small individuals <10 cm).

Decisions regarding the establishment and management of aggregations were primarily taken at the individual level (77.8% of cases), less frequently at the family level (22.2%), and not at the community/tribe level (0%). The selection of aggregation sites was most commonly based on proximity from the respondent’s home (41.9% of cases) and on the previous presence of natural clam populations at the site (34.9%). Less frequently, sites were chosen randomly (11.6%) or based on other geographical or environmental criteria (e.g. proximity to a river mouth). Basic management measures (e.g. removal of predators, repositioning of giant clams after heavy swells or storms, cleaning of algae covering the shells) were eventually applied, but by a minority of owners (29.6%).

Two main collection strategies were observed in relation to restocking the aggregations: some fishers relocated all clams regardless of their size (59.3%), while others (40.7%) only targeted small (i.e. < 10 cm) individuals. Most clam gathering activity was reported to occur during a limited period of 3 to 4 months from June to September (88% of respondents), although a small minority of fishers (12%) also collected clams during the rest of the year (Figure 2b).

The primary motivation for establishing clam aggregations was subsistence consumption (51.9% of responses), primarily for regular household consumption (88.5% of these cases) as opposed to special events (11.5%). Other motivations included the grow-out of juveniles for later consumption (26.9%), and, to a lesser extent, sale (9.6%), recreational purpose (9.6%), and exchange within the community (1.9%) (Figure 3).

**Figure 3.**
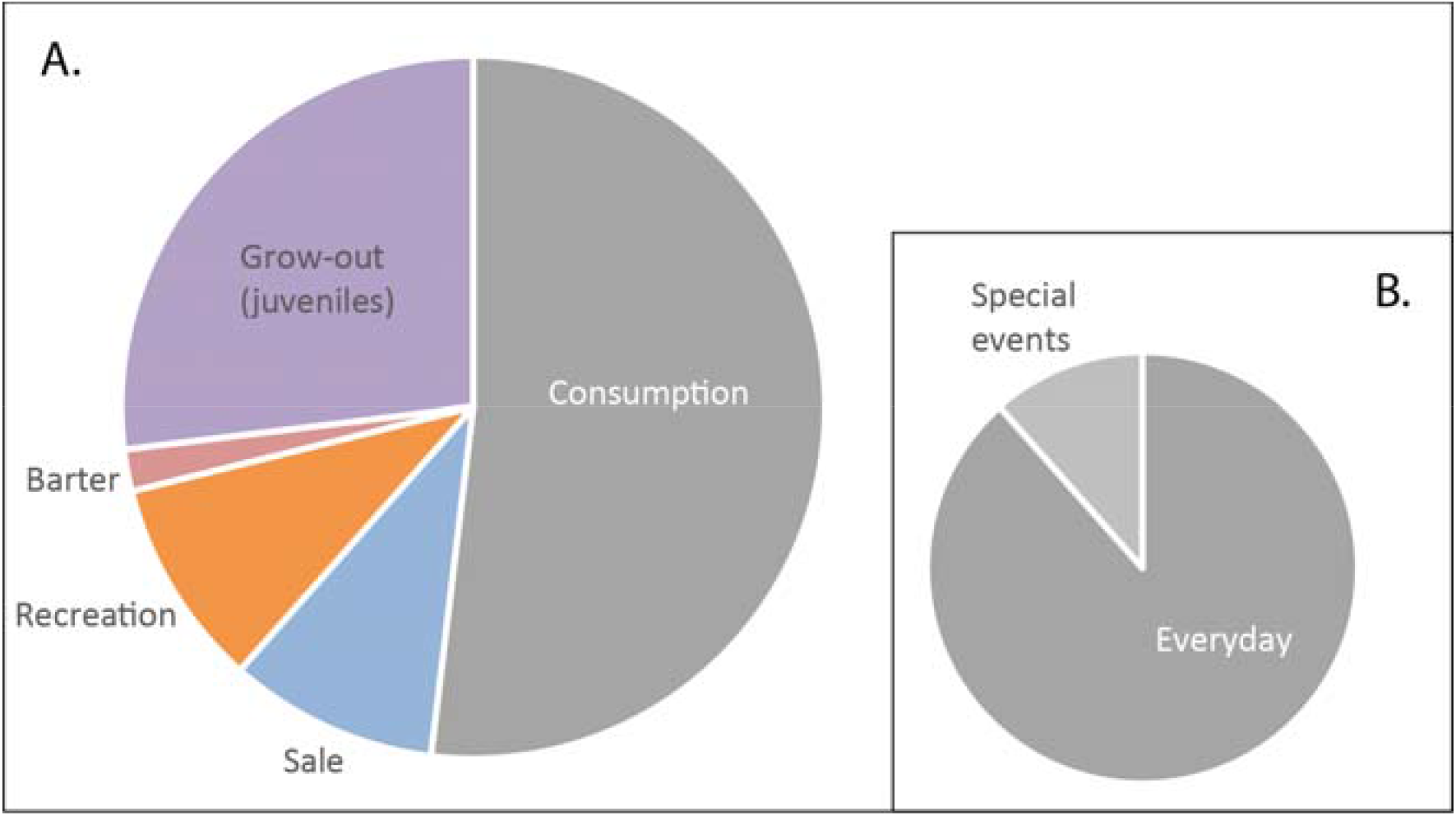
Motivations for creating giant clam aggregations in New Caledonia, South Pacific. A. Percentage distribution of responses given during questionnaire surveys to the question “What is the purpose of clam aggregation?”. B. Distribution of responses for the category Consumption, distinguishing between everyday use and special events.

### 2. Giant clam aggregation surveys

Field surveys conducted along the coastal area of the Tiari tribe identified 17 clam aggregations belonging to 15 people, distributed across the reef flats along a 4.8 km stretch of coastline (Figure 1). The clams were arranged in irregular patches on the substrate, generally delineated by a low wall of coral boulders, and located at varying distances from the shoreline (range: 6 - 310 m). The surface area of the aggregations ranged from 4 to 1 080 m^2^, with most (81%) measuring less than 100 m^2^.

In total, the 17 aggregations contained 7 094 giant clams, with highly variable abundances ranging from 5 to 2 719 individuals per aggregation (mean 417 ± 625 individuals). Two aggregations exceeded 1 000 individuals. Five species were observed in the aggregations: *Hippopus hippopus, Tridacna squamosa, T. maxima, T. derasa*, and *T. crocea*. The assemblages were largely dominated by *Hippopus hippopus*, accounting for 98% of the total abundance (6952 individuals) (Figure 4). In total, 1586 specimens of *Hippopus hippopus* were measured (i.e. 22% of total abundance). Shell length ranged from 6 to 39.5 cm, with aggregations mostly comprising juvenile clams <20 cm (66.7% of assemblages). Young juveniles (≤ 10 cm) and large adults (≥ 30 cm) were virtually absent from the aggregations (11 individuals, 0.7% of the assemblages and 12 individuals, 0.8% of the assemblages, respectively) (Figure 5).

**Figure 4.**
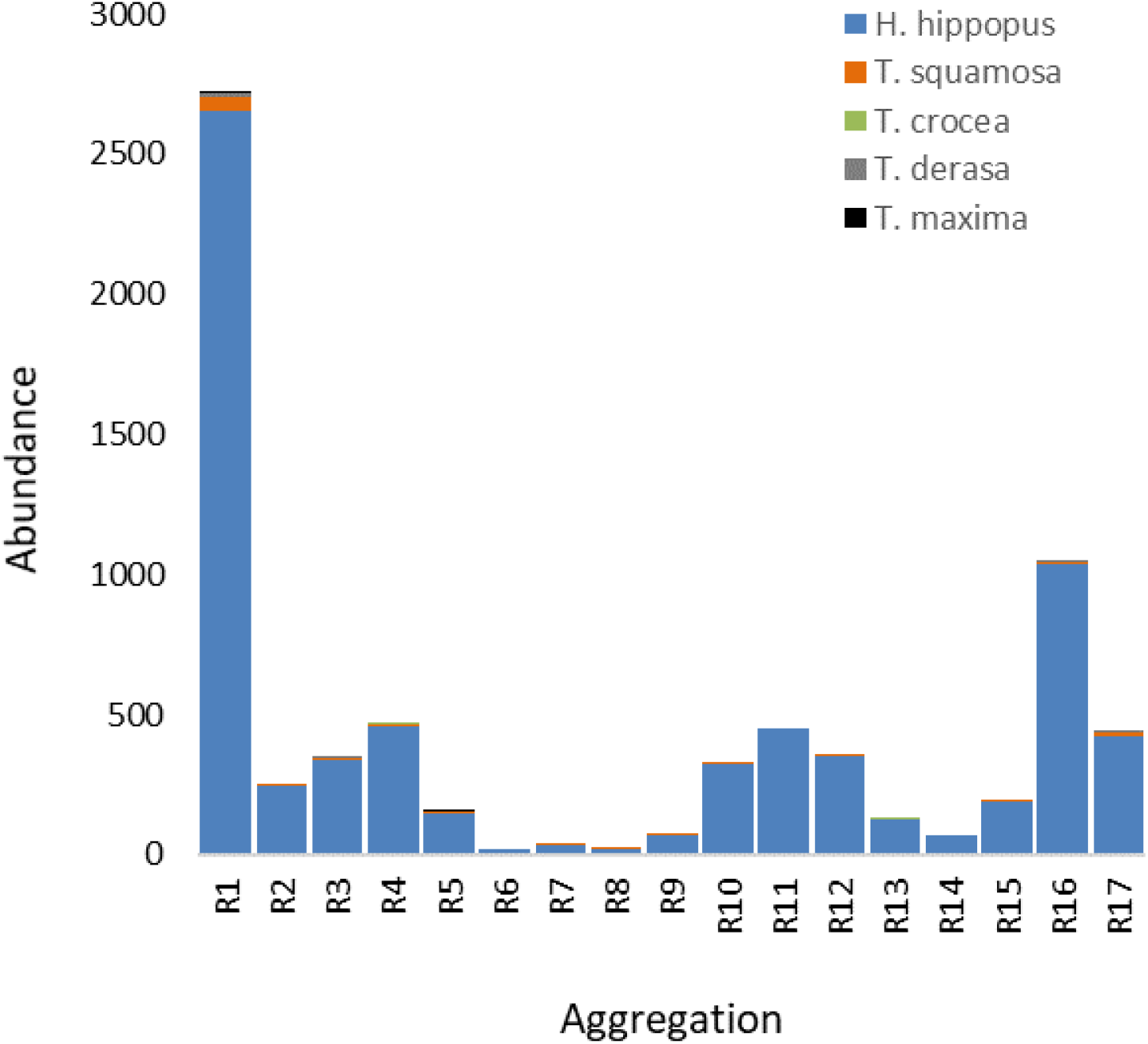
Abundance of giant clams in the aggregations of the Tiari area, North-East coast, New Caledonia based on the aggregation surveys. Total number of individuals per species recorded in the 17 aggregation sites for *Hippopus hippopus, Tridacna squamosal, T. crocea, T. derasa* and *T. maxima*.

**Figure 5.**
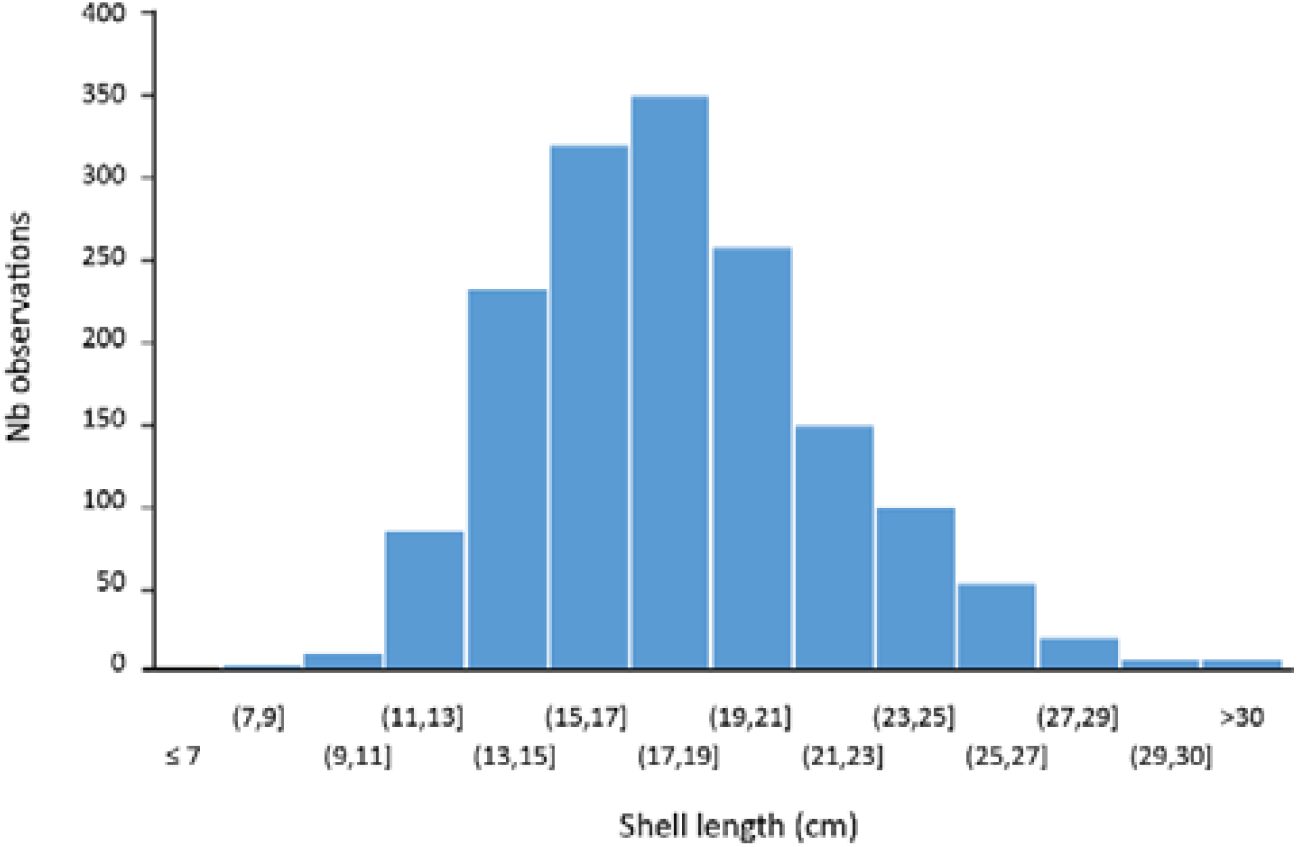
Size distribution of the giant clam *Hippopus hippopus* in the aggregations of the Tiari area, North-East coast, New Caledonia based on the aggregation surveys. Number of individuals per 2-cm shell length class.

### 3. Giant clam population surveys

A total of 120 transects covering an area equivalent to 1 hectare of reef habitat were surveyed along a 3 km stretch of coastline, encompassing Tiari Bay and extending 1 km downstream of the last aggregation site. In total, 282 giant clams belonging to five species were recorded. Abundances were generally low, ranging from 0 to 33 individuals per transect (mean 0.5 ± 0.9 individuals per transect). In contrast to the aggregations, *Tridacna crocea* largely dominated the transects (222 individuals; 78.7% of total abundance), followed by *Hippopus hippopus* (29 individuals; 10.3%), *T. squamosa* (20 individuals; 7.1%), *T. maxima* (10 individuals; 3.5%) and *T. derasa* (a single individual; 0.4%). Average population densities ranged from 1 to 186 clams.ha^-1^ and showed high variability between transects. With the exception of *T. crocea*, observed maximum sizes were significantly smaller than those reported in the literature (Table 2). Although no *H. hippopus* larger than 30 cm was recorded during the survey, the size distribution pattern was broadly similar to that observed in the aggregations: the population was dominated by juveniles <20cm (79.3%), with few reproductive adults (20.7%).

## Discussion

While the harvesting of giant clams is a well-documented practice whose main drivers are generally well understood in most small Pacific Island countries (e.g. Kinch, 2002; Raymakers et al., 2003; Tisdell, 1992), the situation is quite different when it comes to what is referred to in New Caledonia as “giant clam gardens”. To our knowledge, this is the first quantitative study formally addressing these aggregation practices in the region.

### Giant Clam Gardens: comparative framework and history

In New Caledonia, the first reports of clam aggregations date back to the 1970s, mainly as anecdotal observations made in the Northern Province among the East Coast tribes. This practice is believed to have developed in the 2000s, in response to concerns expressed within island and coastal communities about the gradual depletion of the giant clam resource (Tiavouane, 2016). These giant clam gardens should not be confused with the ancient clam gardens described along the north-western Pacific coasts, an early form of mariculture developed by Indigenous communities in North America to increase the productivity of local cold-waters species (Groesbeck et al., 2014; Jackley et al., 2016). Nor are they directly related to restocking programs, such as in Pacific island countries where initiatives have focused on rebuilding natural populations through the grow-out of hatchery-reared juveniles (Bell, 1999; Gomez and Mingoa-Licuanan, 2006). In terms of organization, though not necessarily of objectives, they appear to be more closely related to the “clam circles” described by Chesher (1993) in Tonga, and to the traditional clam gardens initially reported by anthropologists in various locations across the South Pacific including Palau, Papua New Guinea, the Solomon Islands, Kiribati and Tonga (Hviding, 1993; Maclean, 1978; Moir, 1989).

Our findings suggest that, although relatively recent in New Caledonia (one or two generations at most), the practice of “giant clam gardens” is fairly widespread among traditional coastal communities. In total, over 40 giant clam aggregations were documented during our survey of 11 coastal tribes, primarily located in the Northern Province where the majority of the Melanesian population is concentrated (Rivoilan et al., 2021). Additional aggregations were reported by respondents that could not be documented for logistical, environmental or scheduling constraints, including sites along the eastern coast (Houaïlou, Thio, Yaté), the western coast (Poum), and on certain islands such as those in the northern Belep archipelago. The average age of the aggregations identified during this study was 10 years, with only seven aggregations found to be 15 years or older. The oldest aggregation observed in this study was 40 years old, and had been reportedly established by the grandfather of the current owner. We must, of course, keep in mind the potential biases associated with the methodology employed in this study, particularly with regard to the limited spatiotemporal coverage of the surveys and the accuracy of the dates provided by respondents. However, it is unlikely that the existence of much older aggregations would have been systematically omitted during the interviews, especially given the importance and cultural emphasis placed in Melanesian societies on recording and transmitting past events across multiple generations (Hickey, 2006; Ruddle et al., 1992). On the other hand, it is highly likely that the number of aggregations recorded within the surveyed tribes has been underestimated, due to limited access to certain remote areas, the inability to contact some potential owners, and the reluctance (or, in some cases, refusal) of others to share what is perceived as sensitive information.

### Cultural and pragmatic dimensions of giant clam aggregations in New Caledonia

In the Northern Province, giant clam harvesting constitutes a secondary activity, usually conducted in conjunction with fisheries targeting other fish and invertebrate species (Virly, 2004). Our results emphasize that giant clam gardens primarily reflect a pragmatic approach aimed at optimizing access to the resource, rather than an attempt to protect the resource itself. The primary objective, shared by all respondents throughout the study, was to establish a readily available “food reserve,” independent of fishing trips during which clam collection is typically opportunistic rather than planned, and regulated. It is worth noting that this practice somehow exploits a loophole in the provincial regulations, which establish daily harvest quotas per boat for giant clam species. Specifically, non-professional fishers are limited to two clams per boat per day, while professional fishers are allowed up to five clams per boat per day, as stipulated in Article 341-54 of the Environmental Code of the Northern Province and Article 37 of the Environmental Code of the Southern Province. However, no legal limit is specified for gleaning: once giant clams are aggregated on reef flats, these quotas no longer apply, allowing for unlimited collection. In theory, the aggregations, located in the public maritime domain, are accessible to everyone. However, due to the continuity between land and sea, the Kanak people regard coastal areas belonging to their tribe as customary and exclusive property of the tribe or clan (Cavallo et al., 2023). In practice, the system of customary land tenure and traditional access rights prevents outsiders from collecting the clams, which are de facto considered the property of the aggregation owner(s). This observation aligns with the view of Moir (1989) that clam gardens in the South Pacific serve not only to enhance access to the resource, but also to protect appropriated specimens from exploitation by others.

Household consumption was consistently identified by owners as the primary motivation for establishing aggregations, far above income generation or other purposes. These findings are consistent with observations reported for other villages on the east coast of New Caledonia, where approximately 50-60% of non-professional catches were for subsistence consumption, far ahead of sales (30-40%) and of traditional exchange within the community (2-9%) (Faure et al., 2022). Moreover, while certain invertebrate species are particularly sought after in New Caledonia for the preparation of traditional dishes consumed during community events (e.g. lobster *Panulirus* spp., octopus *Octopus cyanea*, trochus topshell *Rochia nilotica* or jumping snail *Conomurex luhuanus*), this does not seem to be the case for giant clams, which are mainly intended for daily household consumption. These findings also align with the estimates provided by Faure et al. (2022), in which giant clams account for up to 2% of daily fishing catches (all fish and invertebrate species combined), but were only targeted for a minority of customary events such as weddings, local fairs or festivals.

### Species-specific composition and management practices in giant clam gardens

In New Caledonia, the practice of giant clam gardens is clearly centered on a single species, *Hippopus hippopus*. It results in quasi-monospecific aggregations in which the presence of other species is either incidental - as in the case of *Tridacna crocea*, which is not actively moved by fishers - or anecdotal, such as *T. squamosa, T. derasa* and *T. maxima*, which are eventually consumed but also regarded as ornamental species by some group owners. The marked preference for *H. hippopus* may be driven by cultural and/or culinary preferences, but more likely by practical considerations. It mostly inhabits very accessible shallow, nearshore habitats and is the only species not anchored to the substrate by a byssus at the adult stage (hence its local name “rouleur”, i.e. “rolling clam”). It can therefore be quickly and easily collected without specific tools, and without damaging the reef matrix. According to the respondents, only adult *Hippopus* measuring between 20 and 30 cm (shell length) are consumed. This size class accounts for only a limited proportion (32.8%) of the individuals measured within the aggregations: in general, fishers tend to avoid collecting individuals that are too large, primarily for logistical reasons such as weight and bulk. Instead, they prefer to target juveniles around 10 cm in size, which are brought back and stocked in the aggregation sites until they reach a consumable size, with the clam gardens functioning as (small-scale) extensive nursery and grow-out areas. The replenishment phase of aggregations generally occurs during spring tides, i.e. the winter months in New Caledonia, when highest tidal range expose large areas of reef flats and gleaning activity is at its maximum (Jimenez et al., 2011).

This study also shows that while notions such as resource protection and conservation were occasionally raised during interviews, they were not explicitly identified as primary motivations for the establishment of giant clam gardens in New Caledonia. This contrasts with community-based practices described elsewhere in the Pacific, where high-value invertebrate species are intentionally relocated and aggregated with the aim of protecting, restoring and/or enhancing their natural populations. Such initiatives were documented for *Hippopus hippopus* and the mother-of-pearl trochus topshell *Rochia nilotica* in small village-based marine reserves (“tabu areas”) in the neighboring archipelago of Vanuatu (Dumas et al., 2010). Similar principles have also been observed in Tonga, where Hviding (1993) noted that, beyond short-term storage for future consumption, “a key motivation for Solomon Islands villagers in taking giant clams from outer reefs to “gardens” off village shore is to protect them from extinction in remote, unsurveilled areas”.

### A practice still unconnected to regional aquaculture approaches

More broadly, in some Pacific countries, in situ giant clam aggregation and grow-out practices have emerged as a natural extension of larger-scale (i.e. supra-community) aquaculture programs, designed to combine conservation objectives with local economic development opportunities. Initiated in the 1970s and 1980s through regional cooperative research efforts, these programs generally involved the establishment of national hatcheries aimed at providing juveniles for restocking projects, often at the community level (Copland and Lucas, 1988; Hart et al., 1999; Heslinga et al., 1984). Starting historically in pilot countries including Palau, Fiji, and the Philippines (among others), these programs were progressively extended to at least twenty Pacific countries, with more than a dozen still reported as active in recent years (Moorhead, 2018; Teitelbaum and Friedman, 2008). This is not the case in New Caledonia, where giant clam aquaculture has not yet succeeded in becoming established, despite trials conducted on *Hippopus* and *Tridacna* spp. in the 1990s, as well as more recent calls for diversification strategies promoted by provincial authorities (Anonymous, 1997; Cluster Maritime Nouvelle-Calédonie, 2016). This is particularly evident in the Northern Province, where *Tridacna* spp. and *Hippopus hippopus* are listed as priority species of strategic interest for aquaculture (Article 232 – Priority aquaculture production, Northern Province Economic Development Code), but only a few private, pilot initiatives have been trialed to date (Cavallo et al., 2023) .

### Potential ecological implications for natural populations

While the practices observed in New Caledonia appear to be driven by more immediate concerns, they may have significant ecological consequences, in particular in terms of population recruitment. The importance of gregarious behaviour for optimal reproduction of benthic macroinvertebrates is now well established (e.g. Ettinger-Epstein et al., 2008; Hadfield and Paul, 2001; Soo and Todd, 2014). Since fertilization success appears to be strongly density-dependent for giant clams, aggregating sparse adults may actually enhance reproduction and subsequent recruitment, an approach historically advocated in various locations across the Pacific to support depleted giant clam population (Lucas, 1994; Teitelbaum and Friedman, 2008). Although few quantitative studies have demonstrated a direct causal relationship, accounts of juvenile clams attributed to the presence of small adult aggregations seem fairly common in the literature. For instance, Chesher (1993) reported sightings of *T. derasa* juveniles only eight months after the establishment of giant clam circles within a community-based marine sanctuary in Tonga. Similar empirical records exist for *H. hippopus* recruits inside clam gardens in the Solomon Islands (Hviding, 1993). Our results showed no significant effect on local recruitment of giant clams, at least at the temporal and spatial scales considered in this study. The relatively high densities of *T. crocea* in Tiari Bay and adjacent reefs (including specimens as small as 1.5 cm) underscore the very limited, if any, fishing pressure on this species, which is generally ignored by local fishers due to its small size and deep position in the reef substrate. Larger species *H. hippopus, T. maxima, T. squamosa* and *T. derasa* exhibited extremely low densities (<= 0.2 individuals per transect), consistent with values commonly reported across the territory for these widely fished species (Dumas et al., 2011; Virly, 2004). Furthermore, no direct evidence of recent recruitment was observed across the 120 transects surveyed in the study area, including up to 1km downstream from the last aggregation. Only four *Hippopus* smaller than 10 cm were found, corresponding to individuals already 2 to 3 years old (Munro, 1992; Pangabean et al., 2018; Shelley, 1989). A similar pattern was consistently observed within all the aggregations where no direct observations of newly recruits could be made, and the proportion of newly recruits could be made, and the proportion of individuals measuring less than 10 cm was negligible.

These findings are not unexpected for organisms with a relatively long pelagic larval duration (7-9 days (Jameson, 1976; Lucas, 1988), which promotes extensive spatial dispersal and generally low self-recruitment rates at the scale of individual reef systems (Kochzius and Nuryanto, 2008; Neo et al., 2013). This was suggested in the Philippines following a massive, long-term restocking program where recruits *T. gigas* were observed from less than a hundred meters up to 20 km from restocking sites (Cabaitan and Conaco, 2017; Requilme et al., 2021). It was further demonstrated in New Caledonia by Tiavouane (2016), who conducted a genetic study on the connectivity of *Hippopus hippopus* populations in the northeast lagoon. The study reported low self-recruitment rates ranging from 0 to 5% across most sampled reefs, while revealing recruitment effects detectable at larger scales, up to at least 35 km from the source population. These results confirm that larval dispersal of giant clams beyond their natal reef is generally predominant; a phenomenon likely amplified by the hydrological and geomorphological features of Tiari Bay, where the openness to the sea and direct exposure to prevailing trade winds are not favorable to the local retention of larvae eventually released from the aggregations. Low levels of local recruitment cannot be entirely ruled out, as suggested by some fishers who occasionally spot juveniles during the winter spring tides from July to August; however direct observation of *H. hippopus* recruits remains particularly difficult due to their small size and greenish-brown coloration, which renders them hard to detect in detritic and seagrass areas.

### Conclusions and perspectives for giant clam management

Their simplicity, the absence of maintenance costs, and their integration into everyday practices, together with a broadly favorable local perception, have contributed to the popularity of giant clam gardens across the Pacific (Anonymous, 2021; Chesher, 1990; Teitelbaum and Friedman, 2008). Understanding whether these traditional practices have a significant effect on recruitment, and perhaps more importantly at which spatial and temporal scales, will require further research effort combining large-scale monitoring with finer-scale (e.g. genetic) approaches. While the direct impact on nearby populations may be limited, their broader importance should not be underestimated, as they definitely contribute to the overall resilience of natural giant clam populations at larger spatial scales, i.e., lagoon and beyond. This particular point would benefit from clearer communication to coastal communities and aggregation owners, in order to better align their expectations within the framework of giant clam restocking projects increasingly considered by local actors, including conservation NGOs, government, and provincial authorities. Given the relatively recent emergence and extent of this practice, as well as the lack of quantitative data on clam turnover within aggregations and on the fishing effort required for their restocking, it remains premature to promote these gardens as a conservation or management tool in New Caledonia. Nonetheless, considering the alarming state of natural stocks and the wish of local communities to continue consuming giant clams, we suggest viewing them as a pragmatic, culturally-embedded approach to local resource use that can provide valuable insights for, though not directly prescribe, management strategies at broader (national) scales.

## Acknowledgements

The authors express their warmest thanks to the technical staff of the IRD Noumea for technical assistance: Miguel Clarque, Gerard Mou-Tham and Joseph Baly. We are also very grateful to the staff of the Northern Province for providing logistical and technical support in the field.

